# Estimating mosquito bionomics parameters with a hierarchical Bayesian model

**DOI:** 10.64898/2026.03.24.713291

**Authors:** Jeanne Lemant, Aurélien Tarroux, Thomas A Smith, Barnabas Zogo, Monica Golumbeanu, Olukayode G. Odufuwa, Seth Irish, Sarah J Moore, Emilie Pothin, Clara Champagne

**Author notes:** Equal contribution (Jeanne Lemant, Aurélien Tarroux).

## Abstract

**Background:** The malaria transmission potential and the vulnerability of *Anopheles* mosquitoes to different vector control methods depend, among other factors, on the endophily, endophagy, anthropophagy and survival of each species. Local information on these bionomic parameters is generally unavailable.

**Methods:** To address this, we estimated species-specific values of these parameters using an augmented version of the global database of bionomics data by Massey et al. (2016). We applied inclusion and exclusion criteria to select eligible studies with relevant experimental designs that minimise bias from collection methods for parous, sac, endophagy, and endophily rates as well as for the resting duration. For the human blood index (HBI), we separated data from indoor and outdoor collections. We fitted hierarchical Bayesian models with levels based on *Anopheles* taxonomy to estimate these quantities. Based on the estimated bionomics, we quantified the expected vectorial capacity reduction after the introduction of a pyrethroid-pyrrole insecticide-treated net (ITN) for 57 *Anopheles* species.

**Results:** We identified 26 eligible studies for endophagy and 61 for the parous rate, leading to a Bayesian posterior average for the *Anopheles* genus of 42% (95% credible interval: 18-70) and 55% (32-77) respectively. HBI values widely varied depending on the location of collection, except for some species showing strong anthropophilic behaviours. Resting duration was estimated to be 2.1 days (1.2 – 4.8) at the genus level. Few studies were available to estimate the sac and endophily rates, which prevented us from deriving precise estimates for the whole *Anopheles* genus. Our estimates of the vectorial capacity reduction following the introduction of a pyrrole-pyrethroid ITN ranged between 48% and 76% across species, highlighting the important differences among mosquito species in vulnerability to vector control interventions.

**Conclusion:** This work demonstrates how data from both *Anopheles* species complexes and individual species can be leveraged to generate species-specific estimates of bionomic parameters, capturing the local characteristics and behaviour of malaria vectors. The dataset is readily updatable as new data become available. However, more frequent and standardised field surveys are still needed to accurately characterise local vector behaviour.

## Introduction

*Anopheles* mosquitoes have long been known to transmit malaria, but not all *Anopheles* are equally competent vectors (Kweyamba et al., 2025; Tadesse et al., 2021). Survival rates of *Anopheles*, and behavioural parameters such as endophily, endophagy, and anthropophagy show considerable spatial and taxonomic variation. These key traits account for much of the geographical variation in malaria transmission intensity (Kiszewski et al., 2004; Macdonald, 1957). Different vectors also differ in their vulnerability to control because vector control interventions, such as insecticide treated nets (ITNs), indoor residual spraying (IRS), or larviciding, target specific behaviours. For example, ITNs and IRS that target indoor biting and resting, have far lower effectiveness in the presence of exophagic and exophilic vectors (Briët et al., 2019; Golumbeanu et al., 2024; Namango et al., 2024; Odero et al., 2024; Sherrard-Smith et al., 2019a; Wang et al., 2024). Transmission models used to inform the choice of the most effective malaria control strategies, tailored to the local context (World Health Organization, 2018, 2025), thus depend on reliable quantitation of the bionomics of local *Anopheles* populations.

Parameterising mathematical models of malaria transmission (e.g. (Chitnis et al., 2008; White et al., 2011)) to measure the impact of vector control interventions at the entomological (Fairbanks et al., 2024; Golumbeanu et al., 2024) or epidemiological level (Briët et al., 2019; Champagne et al., 2025; Sherrard-Smith et al., 2022) has been challenging because of the need to find values for each of the bionomic parameters for each *Anopheles* species present. While some region-specific studies have been conducted (Briët et al., 2019; Wang et al., 2024), it is very demanding to assemble a complete parameterisation even for a location where all the vectors are known.

Field studies of mosquito behaviour are expensive, time-consuming and require specialist expertise. Therefore, we need to make optimal use of existing data, both peer-reviewed and available in the grey literature. A global, systematic collection of available data was made a decade ago (Massey et al., 2016), gathering studies published between 1985 and 2010 from 78 countries. This was accompanied by qualitative summaries of the bionomics of the major vector species (Marianne E. Sinka, Bangs, et al., 2010; Marianne E. Sinka, Rubio-Palis, et al., 2010; Sinka, Bangs, et al., 2011). More than 40 species were identified as dominant malaria vectors (Sinka et al., 2012). While some have been abundantly studied, especially the *An. gambiae* complex in Africa, there are less available data for non-African species. These include locally dominant vectors, such as *An. darlingi* in South America, the *An. sundaicus* complex in Indonesia, or the *Punctulatus* group in Oceania (St Laurent, 2025).

Statistical methods can be used to estimate biometric traits for under-sampled taxa, by using information on the extent to which behaviour tracks phylogenetic relationships. Methods for leveraging phylogenetic relationships to estimate unknown biological traits include phylogenetic imputation using random forest methods (Debastiani et al., 2021), random walk processes on the phylogenetic tree branches (Thorson et al., 2017), or maximum likelihood (James et al., 2021). Hierarchical Bayesian models with different levels for individuals and taxa have been applied to parameterise a model of viral dynamics for different bird species (Banerjee et al., 2017); to estimate traits of an extinct bat species based on measurements of extant relatives (Yohe et al., 2015); to determine whether phylogeny could explain the reaction of *Eucalyptus* tree species to environmental changes (Wooliver et al., 2017), or if the phylogeny of plants could predict demographic rates (Che-Castaldo et al., 2018). In all these cases, data were only available for the lowest level of the hierarchy, corresponding to the tips of the phylogenetic trees, for extant species or individuals. For *Anopheles*, the extent to which behaviour and phylogenetic relationships are linked is unknown. Additionally, challenges arise with datasets such as (Massey et al., 2016): *Anopheles* taxonomy undergoes continual revisions so that older surveys sometimes use different aggregations of species. Moreover, field studies often rely only on morphological identifications, which typically allows classification only to the species complex level, as discrimination of sibling species generally requires molecular methods (Coetzee & Koekemoer, 2013).

In this analysis, we developed six hierarchical Bayesian models, each estimating species-specific values of a bionomics parameter. For each parameter, inclusion and exclusion criteria were defined to select studies from the (Massey et al., 2016) database and the models gave species-specific interval estimates. To account for the complexity arising from identifications of different degrees of precision and for the variation between species in the amount of data, the models borrow information from related taxa, using relevant taxonomic keys and publications (Carnevale & Robert, 2009; Firooziyan et al., 2018; Garros et al., 2005; Harbach, 2004; Marianne E Sinka et al., 2010; Sinka, Rubio-Palis, et al., 2011). An R package `AnophelesBionomics` (https://github.com/SwissTPH/AnophelesBionomics) implementing the Bayesian model is provided to enable users to reproduce the current analysis, or to re-analyse the data using geographical or taxonomic subsets, or their own data. Estimates from the Bayesian models were used to parameterise a model of the full oviposition cycle (Chitnis et al., 2008), which can be applied to estimate vectorial capacity and to model the impact of interventions.

## Methods

The set of parameters to be estimated are those from the model by (Chitnis et al., 2008), as detailed in Table 1.

**Table 1:** Parameters of the mosquito feeding model in (Chitnis et al., 2008) and (Golumbeanu et al., 2024).

| Name | Symbol | Definition | Unit | Bayesian model |
| --- | --- | --- | --- | --- |
| Endophagy | $\beta$ | Preference of mosquitoes for blood-feeding indoors | Percentage | Yes |
| Endophily | $\varphi$ | Proportion of blood-fed mosquitoes resting indoors before searching for a breeding site | Percentage | Yes |
| Human blood index (HBI) (Briët et al., 2019) | $\chi$ | Proportion of resting mosquitoes that have fed on human blood | Percentage | Yes |
| Parous rate (Chitnis et al., 2008) | $M$ | Proportion of host-seeking mosquitoes that have laid eggs at least once | Percentage | Yes |
| Resting period duration (Chitnis et al., 2008) | $\tau$ | Time required for a mosquito that has encountered a host to return to host-seeking (provided that the mosquito survives the whole feeding cycle) | Days | Yes |
| Sac rate (Briët et al., 2019) | $A_0$ | Proportion of parous mosquitoes which have laid eggs the day before. | Percentage | Yes |
| Extrinsic incubation period duration (Chitnis et al., 2008) | $\theta_s$ | Time needed for sporozoites to be present in the salivary glands of a mosquito after biting an infected human | Days | No |
| Oocyst development time (Chitnis et al., 2008) | $\theta_0$ | Time required for oocysts to develop in the mosquito | Days | No |
| Searching time (Chitnis et al., 2008) | $\theta_d$ | Maximum length of time that a mosquito searches for a host in one day if unsuccessful | Days | No |

The analysis was conducted in four steps. Firstly, for each bionomic parameter eligible for inclusion in the Bayesian modelling, we selected studies from (Massey et al., 2016) to ensure that all data resulted from studies conducted under comparable conditions and reporting consistent outcomes. Secondly, we defined genetically related species to inform the taxonomic structure of the Bayesian model hierarchy. Thirdly, we fitted separate hierarchical Bayesian models for each parameter to obtain their respective estimates. Finally, we estimated the reduction in vectorial capacity for different species under an example intervention, a dual active ingredient ITN with chlorfenapyr and alphacypermethrin, to understand how estimates of reduction in vectorial capacity may vary across species and to identify the parameters with the greatest influence on these estimates.

### Selection of relevant datasets and definition of indicators

Each parameter was estimated depending on the availability and variability of the data (Massey et al., 2016). We only retained data collected in the absence of any insecticidal control since its effect is added independently in the model (Chitnis et al., 2008). We also excluded the studies in which authors did not indicate whether some vector control was in place, except when data were very sparse. Data can also be filtered by geographical area of collection to compare estimates in different areas.

Data points were omitted when the sample size was not stated. One observation was defined as a batch of *Anopheles* mosquitoes with a given property. There could be multiple observations per study, if, for example, mosquitoes were collected over several nights or distinguished by species. If the estimated quantity was a percentage, we extracted both the numerator and denominator to account for the sample size. The parameter values were then derived from the number of mosquitoes for which the measured outcome was positive out of those observed.

The six parameters of interest are endophagy, endophily, parous rate, human blood index (indoor and outdoor), sac rate and resting duration. The other parameters required for the mathematical model of (Chitnis et al., 2008) are fixed according to assumptions detailed in a dedicated section below.

Endophagy is defined as the proportion of mosquitoes that bite indoors. We estimated it exclusively using Human Landing Catches (HLC), where adult collectors sit indoors and outdoors with their legs exposed and capture mosquitoes as they land to bite. Endophagy was defined as the number of mosquitoes caught biting indoors, divided by the total number caught biting either indoors or outdoors. HLC generally indicate that mosquitoes are seeking a human blood meal. Comparing HLC counts indoors and outdoors is the standard method to measure endophagy but it tends to underestimate its value in real settings, as it provides mosquitoes with equal opportunity to bite outdoors, whereas most people typically spend most of the night indoors.

Endophily is the proportion of mosquitoes resting indoors after blood-feeding. Pyrethrum Spray Catches (PSC) allow collection of blood-fed mosquitoes resting indoors by spreading white sheets on the floor in the morning and spraying surfaces with insecticide so all resting mosquitoes fall on to the sheets. To also gather data on outdoor-resting mosquitoes, we restricted ourselves to the few studies where window exit traps were placed on windows to collect fed mosquitoes leaving the house to rest outdoors, and PSC were performed in the same houses. Endophily was defined as the ratio between the number of fed mosquitoes collected via spray catches divided by the number of fed mosquitoes collected by both spray catches and window traps. Due to the small number of eligible studies, for this parameter, we included studies in which authors did not indicate whether some control was in place.

The human blood index (HBI) is defined as the proportion of blood meals derived from humans among all the blood meals analysed. It is used to assess whether a mosquito is anthropophilic (high HBI) or zoophilic (low HBI). However, this estimate depends on the collection location: blood-fed mosquitoes found inside houses are more likely to have fed on human beings than those collected outdoors. To account for this potential bias, we separated HBI measured from mosquitoes collected indoors (including window exit traps, since collected mosquitoes were likely feeding indoors) and outdoors and estimated an indoor HBI and outdoor HBI separately. Since the model (Chitnis et al., 2008) requires a single HBI parameter, we relied on indoor HBI only.

To estimate the parous rate, we selected studies reporting the number of parous females as well as the number of inspected mosquitoes. When the percentage of parous females was given together with the number of dissected mosquitoes, we converted it into a number of parous females. The parous rate was defined as the ratio of parous females out of observed females.

The sac rate is the proportion of parous females which have oviposited less than 24 hours before being collected. It is determined by observing the sacs in the ovaries of mosquitoes: if they are distended, the mosquito oviposited recently. The sac rate is the ratio of mosquitoes with distended sacs to overall parous mosquitoes. This quantity is rarely measured and was not considered in (Massey et al., 2016). On the basis of an online search, and after excluding studies performed in the presence of insecticidal control (but retaining those when authors did not indicate whether some control was in place) four eligible publications reporting this quantity were found (Charlwood et al., 2016; Charlwood, Joao Pinto, et al., 2003; Charlwood, J. Pinto, et al., 2003; Molez et al., 1998). The numbers of parous mosquitoes and of mosquitoes with distended sacs reported in these papers were added to the database.

The resting period duration is the period of time blood-fed mosquitoes rest before coming ready to oviposit. It can be estimated either from laboratory observations or from the ratio of fed to gravid females in field collections (Tchuinkam et al., 2010). We could not combine these two types of data together as the field collections are performed under different conditions than in the lab. Because the lab measures did not include the number of mosquitoes observed or the definition of resting period duration (the time until one female becomes gravid, until half are gravid, mean duration for all females…), we decided to rely on field data from the fed/gravid ratio only. If we denote this ratio 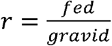, then the resting period duration τ can be approximated by 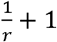. In practice, we estimate *q* = *fed* / (*fed* + *gravid*) using the same hierarchical Bayesian structure as for the other parameters and subsequently derive 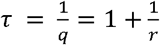. Due to the small number of eligible studies, for this parameter, we included studies in which authors did not indicate whether some control was in place.

### Species classification

A total of 81 distinct species and species complexes appear in the (Massey et al., 2016) database, reflecting the classifications used in the original publications. Some of these names are no longer standard, mostly because the *Anopheles* taxonomy evolves quickly, with species being regularly elevated to complexes. A series of papers (Marianne E. Sinka, Bangs, et al., 2010; Marianne E. Sinka, Rubio-Palis, et al., 2010; Sinka, Bangs, et al., 2011) describe the taxonomy as it stood in the early 2010s. However, additional revisions have occurred more recently, such as the *Punctulatus* complex being now considered as a group (Harbach, 2013).

The first step was to standardise the names. For example, “*Anopheles harrisoni* (formerly *minimus* Sp. C)” was renamed *Anopheles harrisoni* and “*Anopheles nili (ovengensis)*”was standardised to, *Anopheles ovengensis*. Both chromosomal (Savanna, Mopti, Forest, Bamako) and molecular forms (S and M, the latter now recognised as *Anopheles coluzzii*) of *Anopheles gambiae sensu stricto* (*ss*) were reported, although never simultaneously in the same study. Since the chromosomal forms can belong to either molecular form (della Torre et al., 2001), this partial information could be inconsistent so we regrouped all names under *Anopheles gambiae ss/ coluzzii*. Most levels above species were also referred to as complexes, even when the correct or latest terminology should have been subgroup or group. The main example was the “*Anopheles funestus* complex”, which is actually a group containing many species, complexes and subgroups (Harbach, 2013).

We then extracted these names from the whole *Anopheles* genus taxonomy in order to determine the relationships inter-species, inter-levels and between species and the levels above.

The last step consisted of regrouping all the names into categories according to the following criteria:

- There are only two levels in each category (for simplicity of the hierarchical model): the species and a higher level, which for simplicity we call complex but could also be a group or a subgroup.
- A category contains at least two names.
- When we can choose between several levels (group, subgroup and complex), we select the lowest.
- When complex data is ambiguously labelled (e.g. “*Funestus* complex”), it is assumed to refer to the highest category with the same name (in this example “*Funestus* group”). This category is then used as a higher-level category for all the species it contains.

One consequence of these criteria is that we allowed some names to be stand-alone (without any complex).

When classifying the names in the (Massey et al., 2016) database, we always assumed they had been correctly identified, even if not all individual mosquitoes had been identified through PCR, which could mean a species should be reclassified as its complex. Keeping only studies which consistently used PCR for the species identification and reclassifying all the others as the complex would have enormously decreased the total number of available studies.

### Statistical model for parameter estimation across species

We used a hierarchical Bayesian model to infer each bionomic parameter (parous rate, sac rate, endophagy, endophily rates, indoor and outdoor HBI, and resting duration). Each parameter was estimated independently using the same hierarchical structure, comprising three levels: the *Anopheles* genus, the complex, and the species. The distribution of a parameter at a species level is centred around the mean of its complex, whose distribution is itself centred around the mean of the genus (Figure 1).

**Figure 1.**
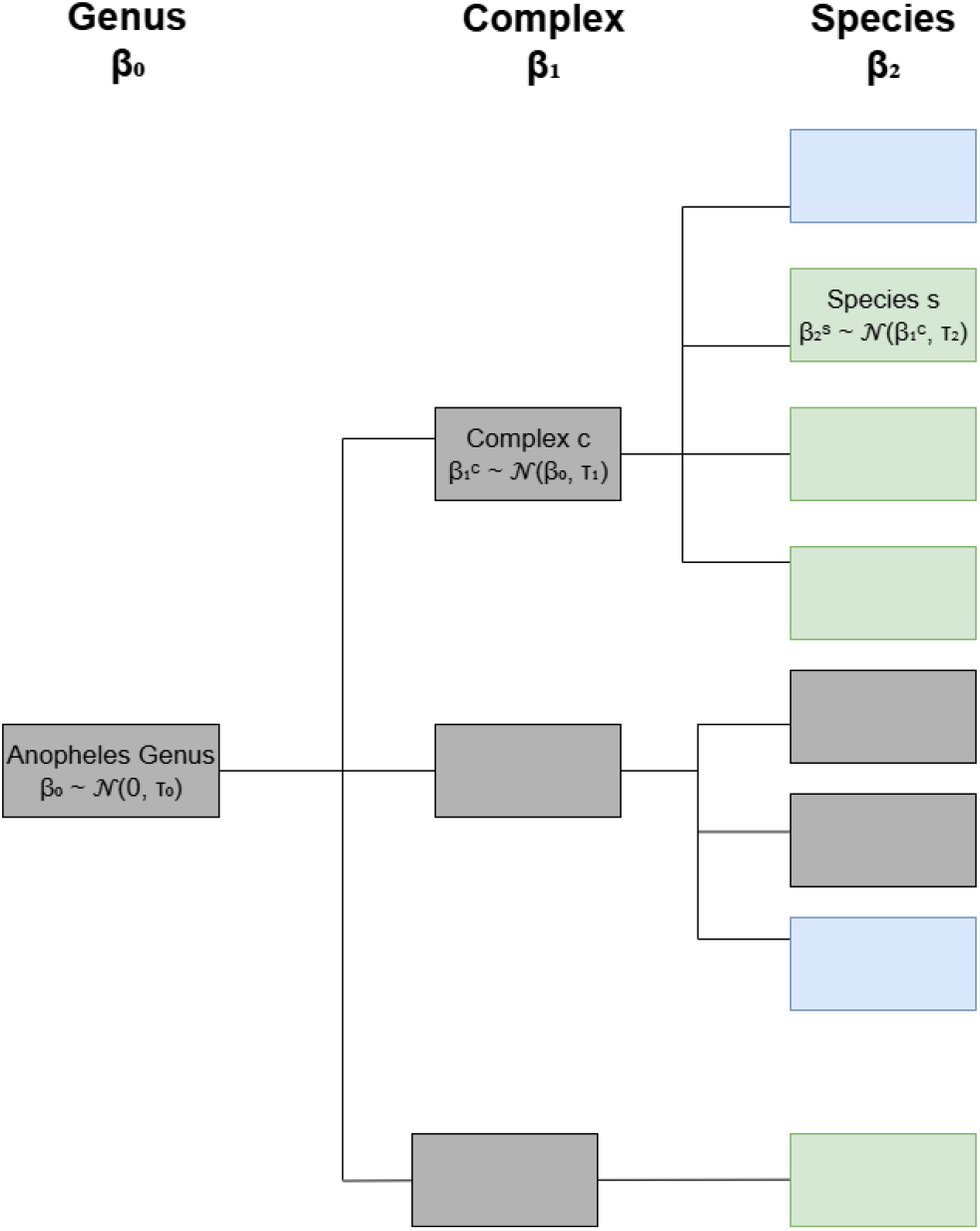
Illustration the hierarchical Bayesian model, with sampled species in green and unlabelled species in blue, and names without any available data in grey. The prior of each level is indicated in the boxes.

The estimated parameters are proportions, so the numbers of mosquitoes which satisfy a property (parous, biting indoors, having ingested human blood,…) were assumed to be binomially distributed. The success probability was the estimated parameter at the species (or complex) level, the sample size the number of observed mosquitoes. The logit of the success probability was equal to the normally distributed priors following the hierarchical structure.

For most data, the species associated with the measurement is provided. However, in some instances, only the complex is reported, without unambiguous identification of the species. Additionally, given the structure of the taxonomy, some complexes do not include any species, and some species are not related to a named complex. Therefore, to ensure consistency and avoid over- or under-weighting certain observations in the hierarchical Bayesian model, we made the following assumptions about the data:

- Species without any complex: we assumed that such species belong to an artificial complex containing only themselves. This prevents them from exerting a disproportionate influence in the genus-level estimation.
- Data for which only complex-level identification is provided:
  - Complex associated with at least one species for which data are available: We assumed that the complex-level information can be represented by a dummy species within the complex, so that all observations carry a comparable statistical weight during estimation. This dummy species is denoted “unlabelled”.
  - Complex with no associated data-available species: We made the same assumption to prevent them from exerting a disproportionate influence in the genus-level estimation.

Let *j* denote each observation, and *s* (*j*) ∈ {1, …, *N*_2_}, the corresponding species (where *N*_*2*_ is the number of sampled species plus the sampled unlabelled species in the initial database). Each species or unlabelled species *s* ∈ [1, …, *N*_*2*_} belongs to a complex *c* ∈ {1,…, *N*_*1*_}, where *N*_1_ is the number of identified sampled complexes in the database.

For a given complex *c, S*_*c*_ is the set of species within this complex.

For complex *c* ∈ 1, …, *N*_1_ and species *s* ∈ 1, …, N_2_, the value of the parameter of interest at the complex level is denoted 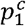 and at species level 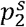

For observation *j* on species 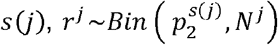, where *r*^*j*^ is the number of mosquitoes with the desired property in observation *j, N*^*j*^ the sample size and 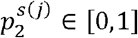 the unknown parameter of species *s* (*j*).

Prior distributions at the different levels (species, complex, genus) were defined on the logit transform of the parameters of interest:

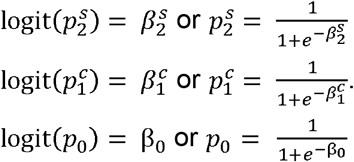

At the overall (*Anopheles* genus) level, β_0_ ∼ 𝒩 0, τ_0_).

For each complex *c* ∈ {1, …, *N*_1_}, 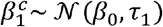, or put otherwise, 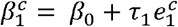, with 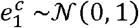 an error term.

For all species *s* ∈ *S*_*s*_ in complex 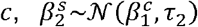, or put otherwise, 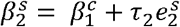, with 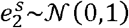 an error term.

The variances of the previously defined normal distributions are given a gaussian hyper-prior: τ_0_ ∼ 𝒩 (0,1), τ_1_ ∼ 𝒩 (0,1), τ_2_ ∼ 𝒩 (0,0.5)

All notations and prior distributions are summarised in Table 2.

**Table 2.** Summary table of the parameters of the Bayesian hierarchical model.

| Symbol | Description | Range/Prior |
| --- | --- | --- |
| **Data** |  |  |
| $j$ | Index for each observation for a species | $\{1, \dots, J_s\}$ |
| $s(j)$ | Species corresponding to observation $j$ | $\{1, \dots, N_2\}$ |
| $r^j$ | Number of mosquitoes with desired property for observation $j$ | $[0, 1]$ |
| $N^j$ | Sample size for observation $j$ | $> 0$ |

| Fitted Parameters |  |  |
| --- | --- | --- |
| $\beta_0$ | $= \text{logit}(p_0)$ | $\mathcal{N}(0, \tau_0)$ |
| $\beta_1^c$ | $= \text{logit}(p_1^c)$ | $\mathcal{N}(\beta_0, \tau_1)$ |
| $\beta_2^s$ | $= \text{logit}(p_2^s)$ | $\mathcal{N}(\beta_1^c, \tau_2)$, |
| $\tau_0$ | Variance for *Anopheles* genus level | $\mathcal{N}(0,1)$ |
| $\tau_1$ | Variance for complex-level variation | $\mathcal{N}(0,1)$ |
| $\tau_2$ | Variance for species-level variation | $\mathcal{N}(0,0.5)$ |
| Transformed quantities |  |  |
| $p_0$ | Genus-level parameter | $[0,1]$ |
| $p_1^c$ | Parameter for complex $c$ | $[0,1]$ |
| $p_2^s$ | Parameter for species $s$ | $[0,1]$ |

The model was fitted using Stan (Carpenter et al. 2017), via the RStan package (Stan Development Team 2024) with four chains of 3000 iterations, including a burn-in of 1500 iterations. Convergence was assessed using the R-hat diagnostics provided by RStan.

For each of the estimated parameters, we obtained the Bayesian posterior distribution of the parameter for each sampled species and complex. We then borrowed the estimates of a complex for a non-sampled species within this complex. If no data were available at the complex level either, we used the estimate for the *Anopheles* genus, which summarises all available data. To estimate the vector model parameters, we extracted the empirical mean of the posterior distribution. The posteriors distributions are as follows:

- 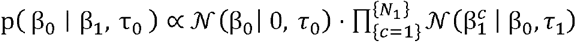
- 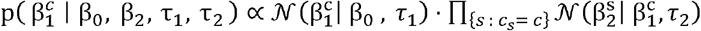
- 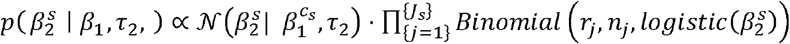
- *P* (τ_0_ | β) ∝ 𝒩 (τ_0_ | 0,1) · 𝒩 (β_0_ | 0, τ_1_)

Once the posterior distributions were obtained, highest posterior density (HPD) regions were computed for each parameter using the coda package (Plummer et al. 2006). For each chain generated by Stan, the HPD interval was determined at the 99% credibility level, and samples lying outside the corresponding HPD region were discarded. This procedure ensures that subsequent summaries and comparisons are based exclusively on the most credible portion of the posterior distribution.

We developed the R package `AnophelesBionomics` (https://github.com/SwissTPH/AnophelesBionomics) to implement this framework and facilitate reproducible analyses. Users can estimate bionomic parameters using the provided database or their own data, with filtering options by geographic region and time period. When constructing the hierarchical Bayesian model, the package enables users to specify the number of chains, burn-in iterations, and total iterations, providing flexibility to tailor model fitting to specific convergence diagnostics and computational resources.

Once the model is fitted, the package returns posterior estimates at the species, complex, and genus levels, and includes functions for visualizing posterior distributions and summary statistics.

### Vectorial capacity calculation

Vectorial capacity is defined as the total number of potentially infectious bites originating from all the mosquitoes biting a single perfectly infectious (i.e. all mosquito bites result in infection) human on a single day. It represents the potential for a given mosquito population to transmit malaria (Garrett-Jones and Grab 1964). Vectorial capacity is calculated using the approach by (Chitnis et al. 2008), as implemented in the AnophelesModel R package (Golumbeanu et al., 2024).

We calculated vectorial capacity in the presence and in the absence of a vector control intervention, taking the example of a chlorfenapyr-alphacypermethrin ITN as estimated from the study by (Odufuwa, in prep) in (Champagne et al., 2025), with 80% use.

To account for ITN effectiveness decay, the model calculates the steady-state vectorial capacity over a 3-year period, updating effective coverage and entomological efficacy at each time step based on functional survival and insecticidal durability. The reductions in vectorial capacity are averaged to provide a summary value for the 3-year period. Functional survival estimation follows the methodology of (Champagne et al., 2025), using a Weibull decay function fitted to observed attrition data from (Martin et al., 2024). The effective half-life and decay shape parameter are estimated by minimizing the least squares distance between the data and the model.

For calculating human in-bed exposure, we use data from the AnophelesModel R package (Golumbeanu et al., 2024). First, we compute the hourly averages for the biting proportions indoors and outdoors, as well as the proportion of humans in bed and indoors. For each species with available data, hourly averages are computed. If no data are available for a given species, the calculation is performed at the complex level: when at least one species within the complex has data, the complex value is taken as the mean across all available species. If no species within a complex has data, the corresponding genus value is used instead, defined as the mean across all complexes within that genus. The resulting average activity patterns for each species are provided as Supplementary Information. Once these activity averages are determined for both mosquitoes and humans, we use the same AnophelesModel package to calculate the exposure levels. Exposure coefficients calculated using the methodology by (Golumbeanu et al., 2024) are multiplied with the ITN use value, as in (Champagne et al., 2025). All input parameters are summarized in Supplementary Table 2.

The model uses as inputs the estimates from the Bayesian hierarchical models for the parous rate, the sac rate, and the overall human blood index. The overall human blood index is defined as the average between the indoor and the outdoor HBI, weighted by endophagy. The other parameters listed in Table 1 were fixed as follows:

**The searching time** is always multiplied by relative availability in the feeding cycle model (Chitnis et al., 2008) and is defined for homogeneity of units, so there is no need to estimate it or to let it vary by species. We fix its value at 0.33 day, as in previous literature (Briët et al., 2019).

**The extrinsic incubation period duration** is the time needed for sporozoites to be present in the salivary glands of a mosquito after biting an infected human. It is mostly temperature-dependent so we applied the Detinova degree-day model (Ohm et al., 2018)

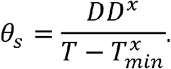

This model assumes that a *Plasmodium* parasite *x* (*falciparum* or *vivax*) needs a certain number of days where the mean temperature *τ*_*mean*_ is above the minimal temperature suitable for its development of parasite 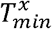. A degree-day is the difference between the mean daily and the minimal suitable temperature, and *DD*^*x*^ is the sum of degree-days required for the sporozoites to mature. For *P. falciparum, DD* = 111 and *τ*_*min*_ = 16°*C*, whether for *P. vivax, DD* = 105 and *τ*_*min*_ = 14.5°*C* (Ohm et al., 2018).

We set *θ*_*s*_ = 10 days as a default value, which corresponds to a temperature of 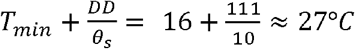 for *P. falciparum*, and 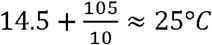 for *P. vivax*.

**The oocyst development time** is the time between ingestion of male and female gametocytes present in human blood and presence of oocysts in the mosquito midgut. It mostly depends on temperature and was fixed to 5 days as in (Chitnis et al., 2008).

Vectorial capacity reduction was calculated for each species by taking 200 samples from the posterior distributions of each parameter. Mean and 95% credible intervals over these samples were reported.

Additionally, we conducted a sensitivity analysis on the contribution of each input parameter (parous rate, sac rate, indoor and outdoor HBI, endophagy, resting duration) to the vectorial capacity reduction associated with the deployment of the example ITN. A random sample of 20000 values was drawn from the posterior distribution for each parameter of the *Anopheles* genus. Partial rank correlation coefficients (PRCC) were computed to assess the relative contribution of each input to the vectorial capacity (Iooss and Lemaître 2015). This was implemented using the sensitivity R package (Iooss et al. 2020). Endophily was not included as this parameter only affects the impact of interventions targeting resting mosquitoes, such as IRS (Golumbeanu et al., 2024).

## Results

### Taxonomy

Putting together the standardised names from the database and the sac rate studies, we obtained 57 species and 17 complex, subgroup or group names (Figure 2).

**Figure 2.**
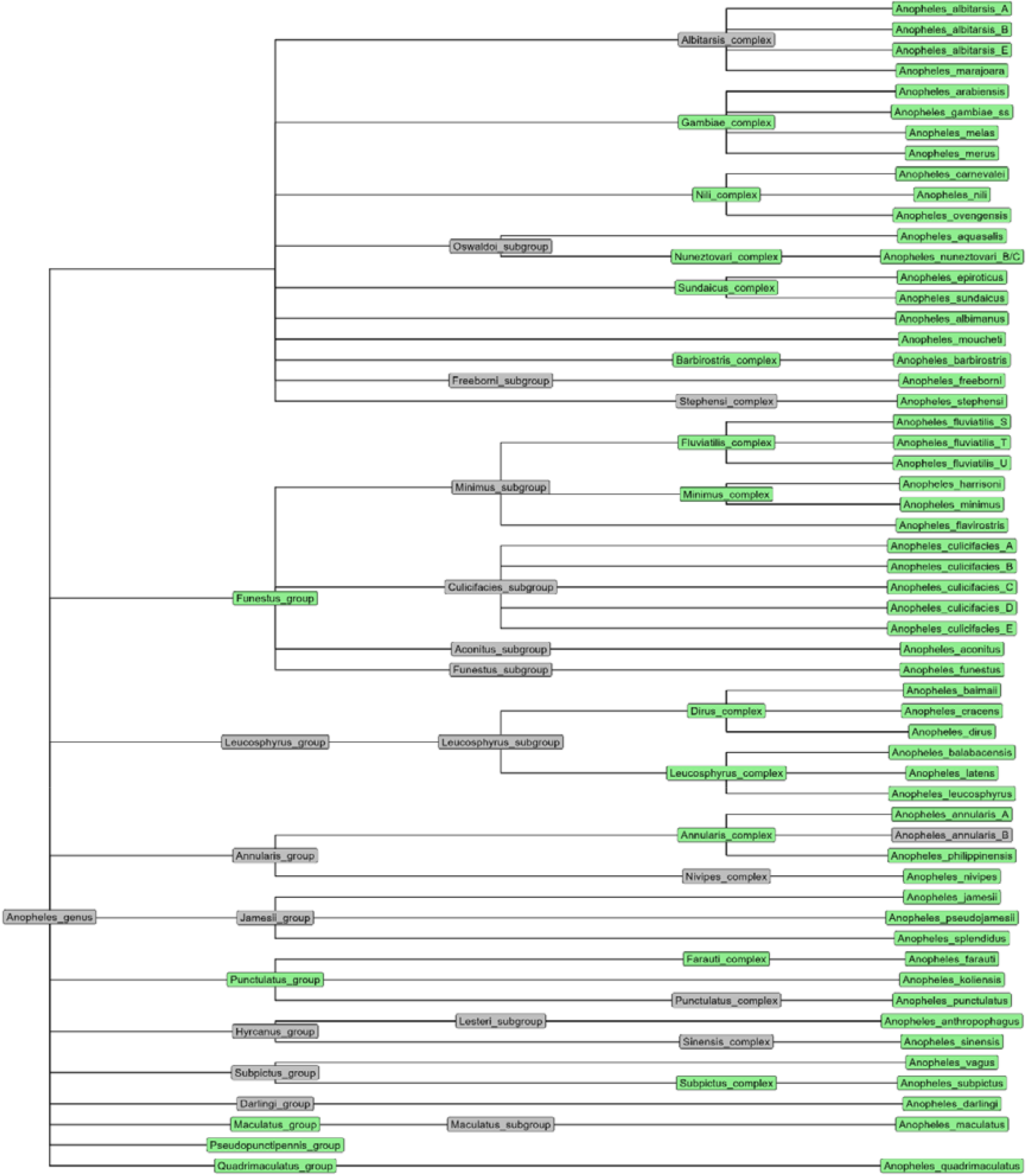
Subset of the Anopheles genus taxonomy with names sampled in the (Massey et al., 2016) database in green and those absent from the database in grey.

The obtained categories are listed in Supplementary Table 1, together with the stand-alone names.

### Selected data

Flow charts describing the application of inclusion and exclusion criteria for all indicators are available in Supplementary material. We selected most data on parous rate (521 observations on 11 species and 11 complexes from 61 studies) and least data on endophily (six observations on two species and two complexes from four studies). In total, 12 complexes and 21 species had data for at least one parameter, but data were sparse, with at most 11 complexes and 11 species sampled for the same parameter (parous rate). Sample sizes ranged from one to 15,467 observed mosquitoes (for the parous rate).

For some parameters such as the parous rate, we have numerous observations from different complexes and species, and relatively consistent data (standard deviation 0.21). Endophily is the parameter with the fewest data points and the most dispersed values (standard deviation 0.35) (Table 3).

**Table 3.** Summary of available data for each parameter across surveyed mosquito species and complexes. The table includes the number of surveys, observations, standard deviations of the observed value of the parameters, and the number of species and complexes observed for each parameter

|  | Parous rate | Endophagy | Endophily | Sac rate | Indoor HBI | Outdoor HBI | Resting duration |
| --- | --- | --- | --- | --- | --- | --- | --- |
| **Number of observations** | 541 | 223 | 8 | 35 | 224 | 78 | 59 |
| **Sample size** | 196,954 | 89,219 | 1,419 | 4,327 | 38,583 | 4,739 | 18,646 |
| **Standard deviation of the observed value** | 0.21 | 0.24 | 0.35 | 0.28 | 0.27 | 0.28 | 5.98 |
| **Number of species** | 11 | 7 | 2 | 10 | 7 | 5 | 9 |
| **Number of complexes** | 11 | 6 | 2 | 5 | 3 | 3 | 7 |

There was only one study on sac rate performed without any insecticide control, with 1,396 observed mosquitoes and a sac rate of 40%. We therefore also included three studies for which the insecticide control had not been reported, resulting in 35 observations for 10 species and 5 complexes (Table 3).

For all parameters, except for outdoor HBI, a large proportion of mosquitoes analysed belonged to the *Anopheles gambiae* complex, ranging from 27% for the sac rate up to 82% for endophily. The *Anopheles funestus* complex was also well represented across all parameters (Figure 3).

**Figure 3.**
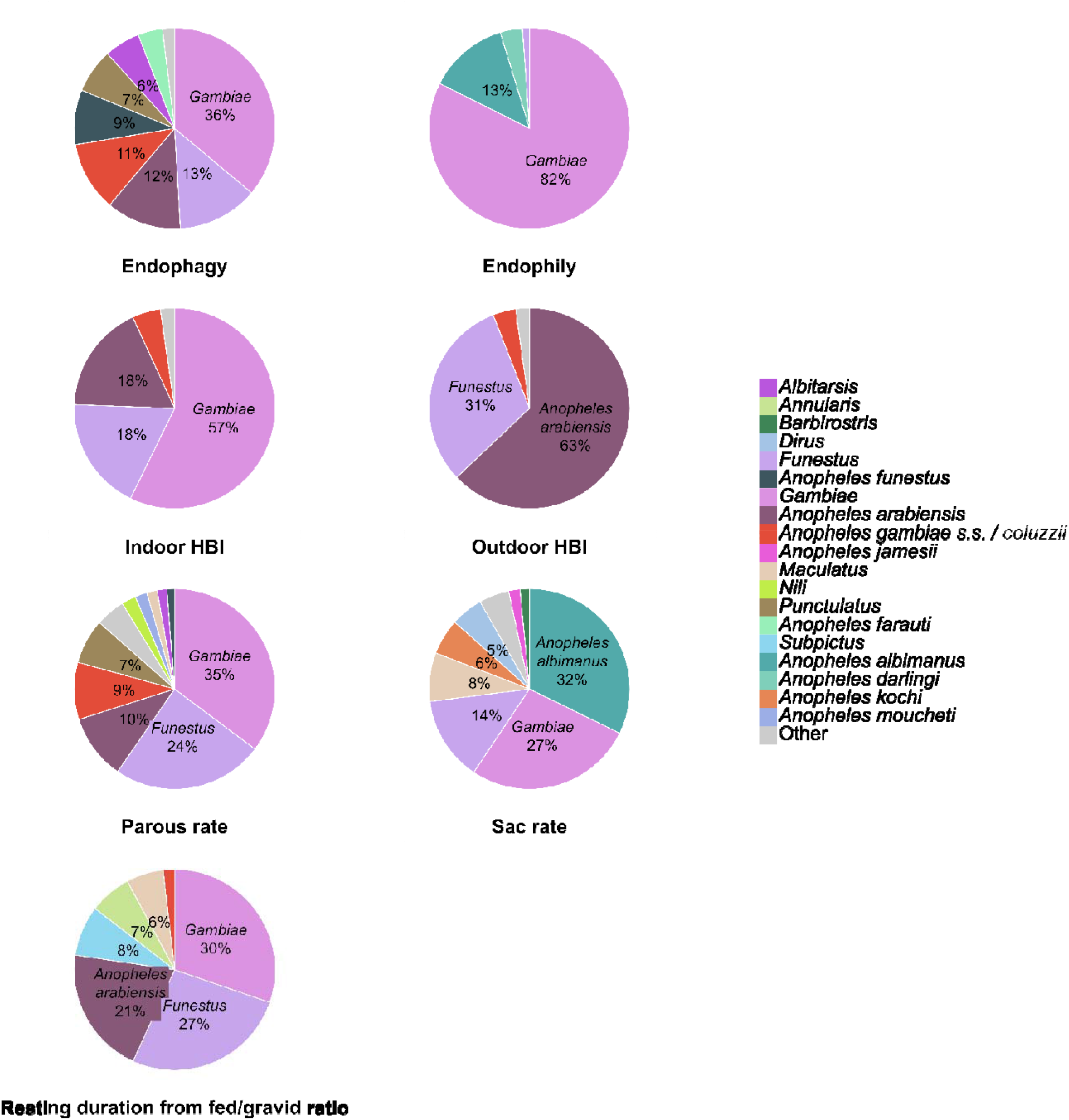
Percentages of observations in the dataset categorized by species or complex for each parameter. Each slice corresponds to the percentage of data points associated with a given group (slices below 5% not labelled, slices above 20% include species/complex names). In the legend, “Complex” refers to data for which identification was available at the complex level only.

For endophagy, parous rate, resting duration and sac rate, observations were distributed across a wide range of species and complexes, such that no single species or complex dominated the dataset. In contrast, for endophily, outdoor HBI, and indoor HBI, a single species or complex accounted for most observations (Figure 3).

### Hierarchical Bayesian model

In this section, for readability, only the posterior distributions of a subset of species and complexes are shown for some parameters. The full figures are provided in the Supplementary material (Supplementary Figures 3 to 15) and the full table with each species’ estimates as Supplementary information. Additionally, the supplementary material provides a comparison of the Bayesian estimates with the raw weighted average for each parameter, for species in the *Gambiae* and *Funestus* complexes/groups as an example (Supplementary Figures 16 and 17).

### Endophagy

Based on paired indoor-outdoor HLC, mean endophagy weighted by sample size was 45 ± 24%, while the estimated endophagy for the whole *Anopheles* genus was 42% (95% credible interval: 18-70). We found that on average 29% (18-46) of *Anopheles albimanus* are endophagic, while the endophagic rate was 53% (52-54) for *Anopheles funestus*, 56% (55-57) for *Anopheles gambiae sensu stricto* / *coluzzii* and 66% (65-67) for *Anopheles arabiensis* (Figure 4).

**Figure 4.**
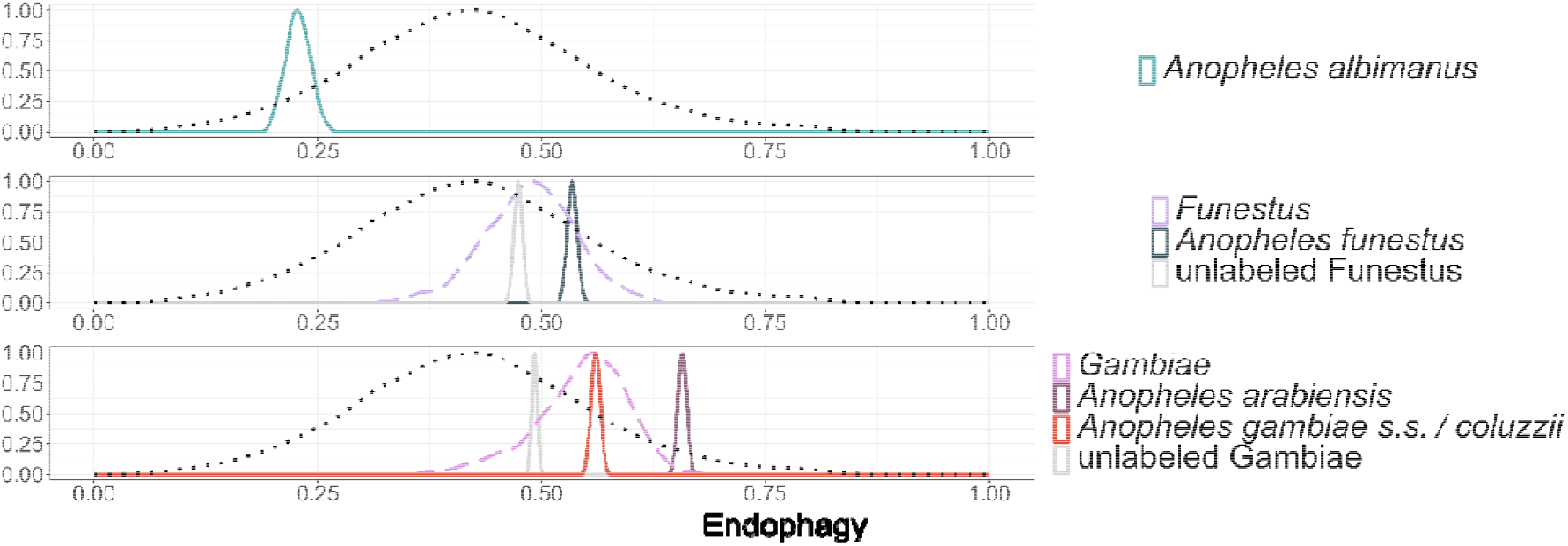
Posterior densities for endophagy for a subset of species. The black dotted line is the posterior density for the pooled estimate at the genus level, the coloured dashed lines are for each complex, and the coloured solid lines for individual species. Y-axis is normalised to ease visualisation of all curves. The full figure including all species is provided in the Supplementary material (Supplementary Figure 3).

When separating countries between East and West Africa, we observed some marked differences for *Anopheles arabiensis* (67% (66 - 68) in East Africa, 38% (33 - 43) in West Africa) and *Anopheles gambiae sensu stricto / coluzzii* (26% (21 - 32) in East Africa, 57% (56 - 58) in West Africa), while *Anopheles funestus* endophagy was more similar between the two regions (respectively 46% (26 - 66) and 53% (52 - 55)) (Supplementary Figure 4 and Supplementary Figure 5).

### Endophily

We found that 69% (36-91) of *Funestus* complex mosquitoes were endophilic, 74% (46-88) of *Gambiae* complex and 45% (2-97) of the whole *Anopheles genus* (Figure 5). Despite the small sample size (51 mosquitoes) of the only observation for *Anopheles darlingi* (Supplementary Figure 7), the posterior density was still very precise since the observed endophily was 2% (0-7) and the values were restricted to the interval by the logit transform.

**Figure 5.**
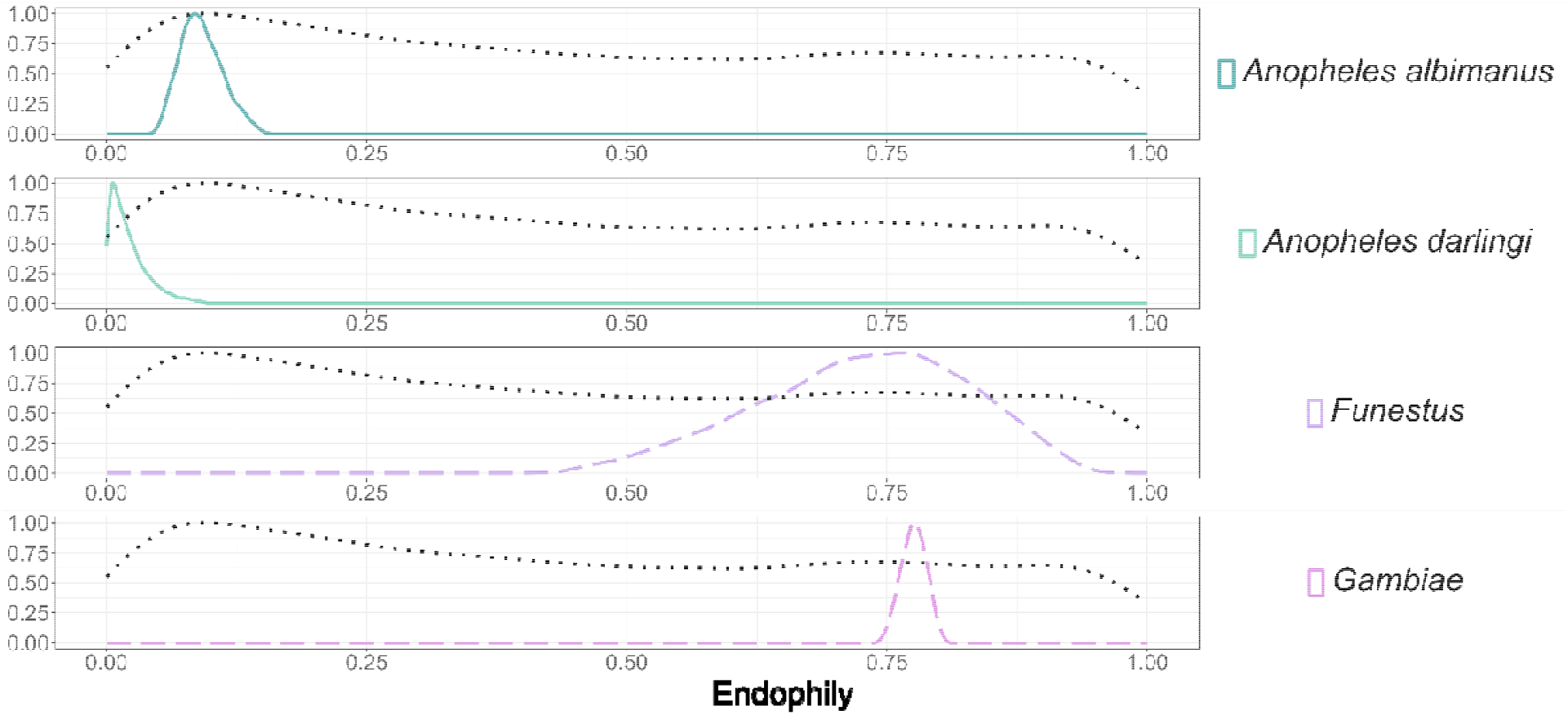
Posterior densities for endophily. The black dotted line is the posterior density for the pooled estimate, the coloured dashed lines, for each complex, and the coloured solid lines for individual species. Y-axis is normalised to ease visualisation of all curves.

The presence of both endophilic and exophilic species within the *Anopheles* genus leads to high variability in the estimates of endophily at the genus level. The hierarchical model captures this heterogeneity with a wide standard deviation and a flat distribution at the genus level. The lower peak in the distribution is driven by observations from *Anopheles albimanus* and *Anopheles darlingi*, which together represent 17% of all genus-level data (Figure 3), and the rest of the distributions driven by observations from the *Funestus* group and *Gambiae* complex, which represent the rest of the observations.

### Human blood index

The posterior densities illustrate how the behaviour of *Anopheles arabiensis* varies depending on location and is likely driven by host availability, with a mean HBI of 65% (64-66) from indoor collections and 13% (12-15) from outdoor collections, while *Anopheles gambiae sensu stricto / coluzzii* feeds mostly on human beings: its mean posterior density was 86% (85-88) from indoor collections and 70% (64-77) from outdoor collections (Figure 6 and Figure 7).

**Figure 6.**
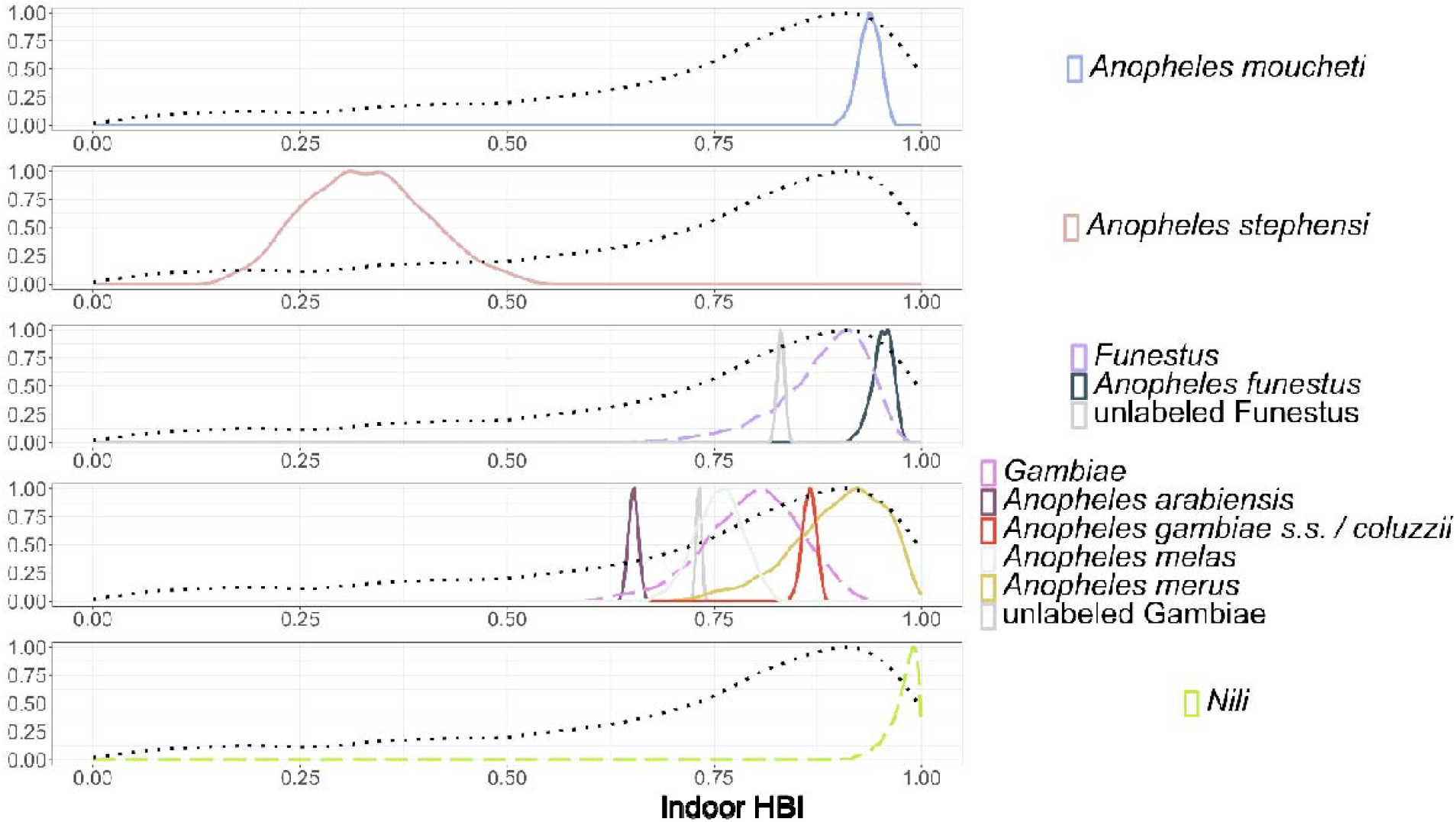
Posterior densities for indoor HBI. The black dotted line is the posterior density for the pooled estimate, the coloured dashed lines, for each complex, and the coloured solid lines for individual species. Y-axis is normalised to ease visualisation of all curves.

**Figure 7.**
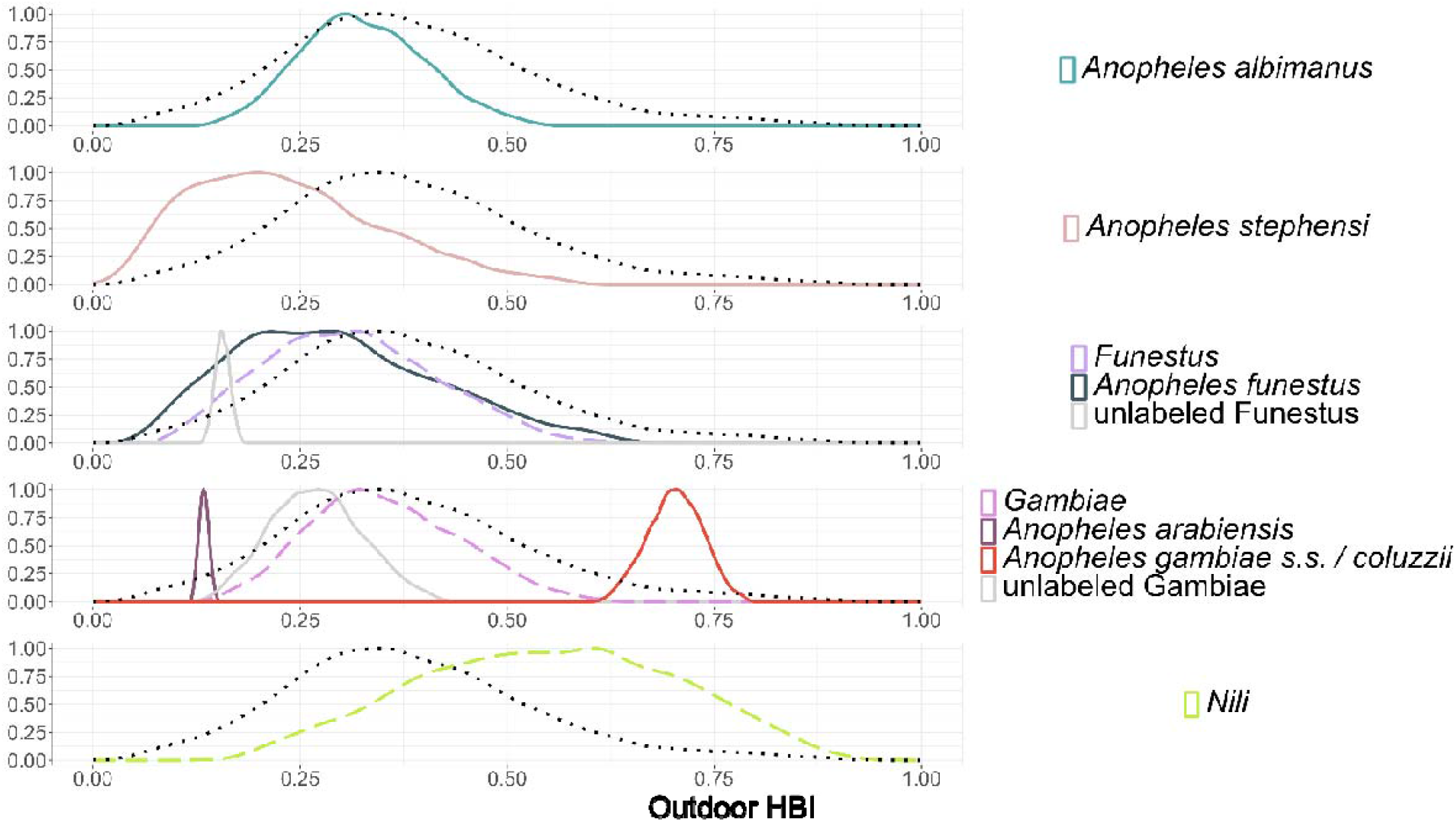
Posterior densities for outdoor HBI. The black dotted line is the posterior density for the pooled estimate, the coloured dashed lines, for each complex, and the coloured solid lines for individual species. Y-axis is normalised to ease visualisation of all curves.

The large differences in biting preferences across species cause the estimates of both indoor and outdoor HBI for the whole *Anopheles* genus to exhibit a flattened distribution. This results in high uncertainty around the *Anopheles* genus estimates: 73% (12-99) for indoor HBI and 38% (12-73) for outdoor HBI.

### Parous rate

The weighted average of the selected data for the parous rate was 66%, while the estimated parous rate for the whole *Anopheles* genus was 55% (32-77). This highlights the shrinkage effect in the Bayesian models, through which the effects of sampling variations are reduced (cf. Supplementary Figures 16 and 17 for further examples on the *Gambiae* and *Funestus* complexes/groups). Overall, there was a lot of variability within the parous rates, with values ranging from 35% (25-50) for the *Nuneztovari* complex to 76% (74-77) for *Anopheles funestus sensu stricto* (Figure 8).

**Figure 8.**
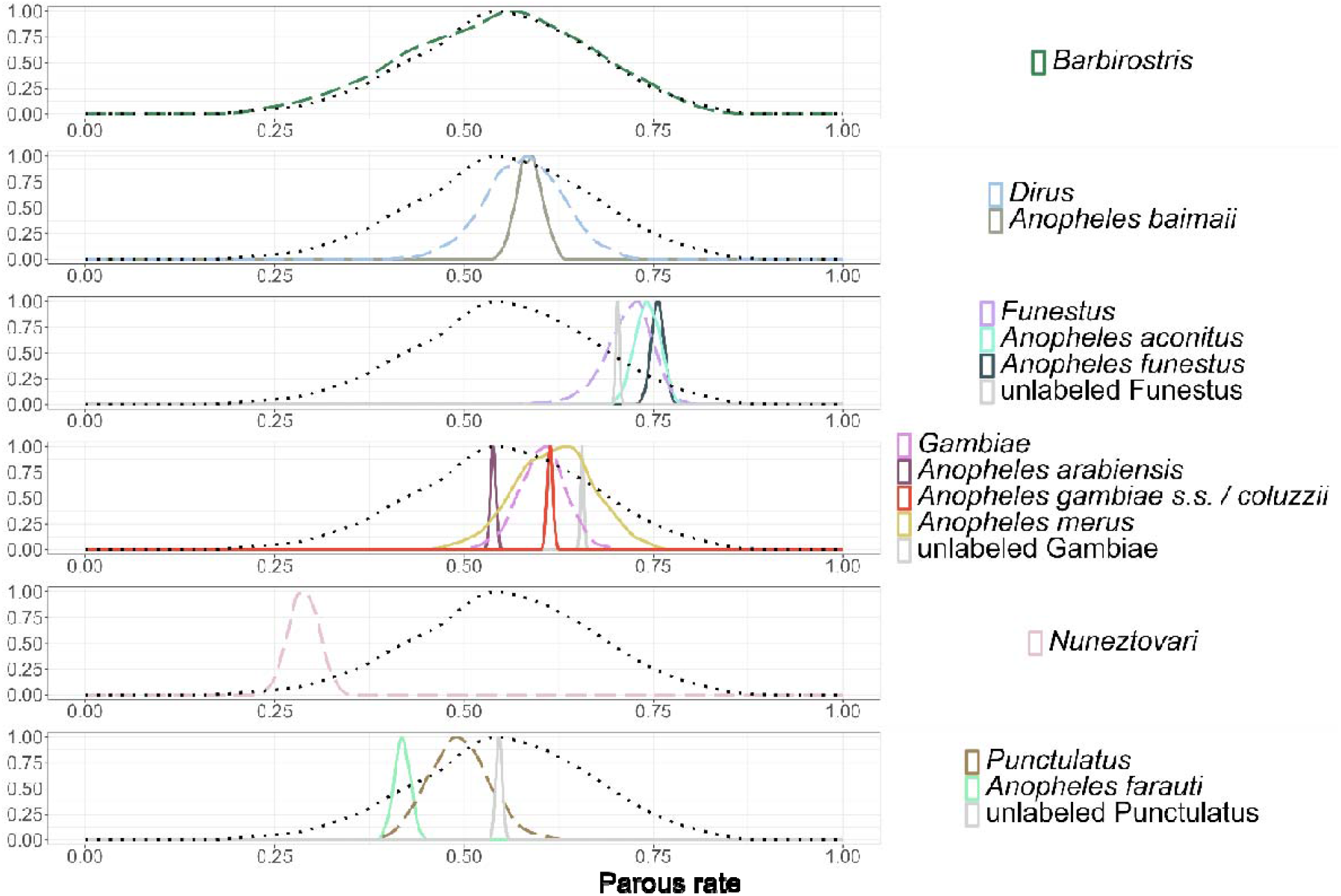
Posterior densities for the parous rate for a subset of species. The black dotted line is the posterior density for the pooled estimate, the coloured dashed lines, for each complex, and the coloured solid lines for individual species. Y-axis is normalised to ease visualisation of all curves. The full figure including all species is provided in the Supplementary material (Supplementary Figure 12).

In the case of *Anopheles merus* there were only three observations with small sample sizes observations (1, 13 and 15 mosquitoes), so its estimate lies within the complex, with a mean of 62% (52-72) (Supplementary Figure 12).

We obtained posterior densities for *Dirus* complex and *Hyrcanus* group even if they were not sampled, since species within these complexes were still sampled. This is the reason why the *Anopheles baimaii* estimate lies within the *Dirus* complex estimate: only *Anopheles baimaii* was sampled, and the *Dirus* posterior is therefore a compromise between the species and the genus (Figure 8).

The posterior estimate for *Anopheles barbirostris* aligns with that of the *Anopheles* genus primarily because there are insufficient data at the species level. In fact, for *Anopheles barbirostris* we have only a single observation with a sample size of two (Supplementary Figure 11).

For the *Punctulatus* complex, however, we had data at both the *Punctulatus* complex and *Anopheles farauti* species levels. Here, the complex serves as a perfect compromise between the posterior densities of the unlabelled *Punctulatus* complex and *Anopheles farauti* (Figure 8), despite the fact that the sample sizes differ by an order of magnitude: 842 and 1218 *Anopheles farauti* mosquitoes were observed, whereas 9464 and 4779 were counted as *Punctulatus complex* (Supplementary Figure 11). As a result, the smaller sample size for *A. farauti* leads to greater uncertainty in its posterior estimates.

### Resting duration

The posterior distributions of the resting duration (in days) revealed clear differences between species ( Figure 9). The *Subpictus* complex showed the highest resting duration, with a resting duration of 2.2 days (1.4-4.0). In contrast, *Anopheles gambiae sensu stricto* / *coluzzii* exhibited shorter resting times with an estimation of 1.1 days (1.0-1.1), while the *Funestus* group displayed intermediate values close to 2.0 days (1.3-3.4).

**Figure 9.**
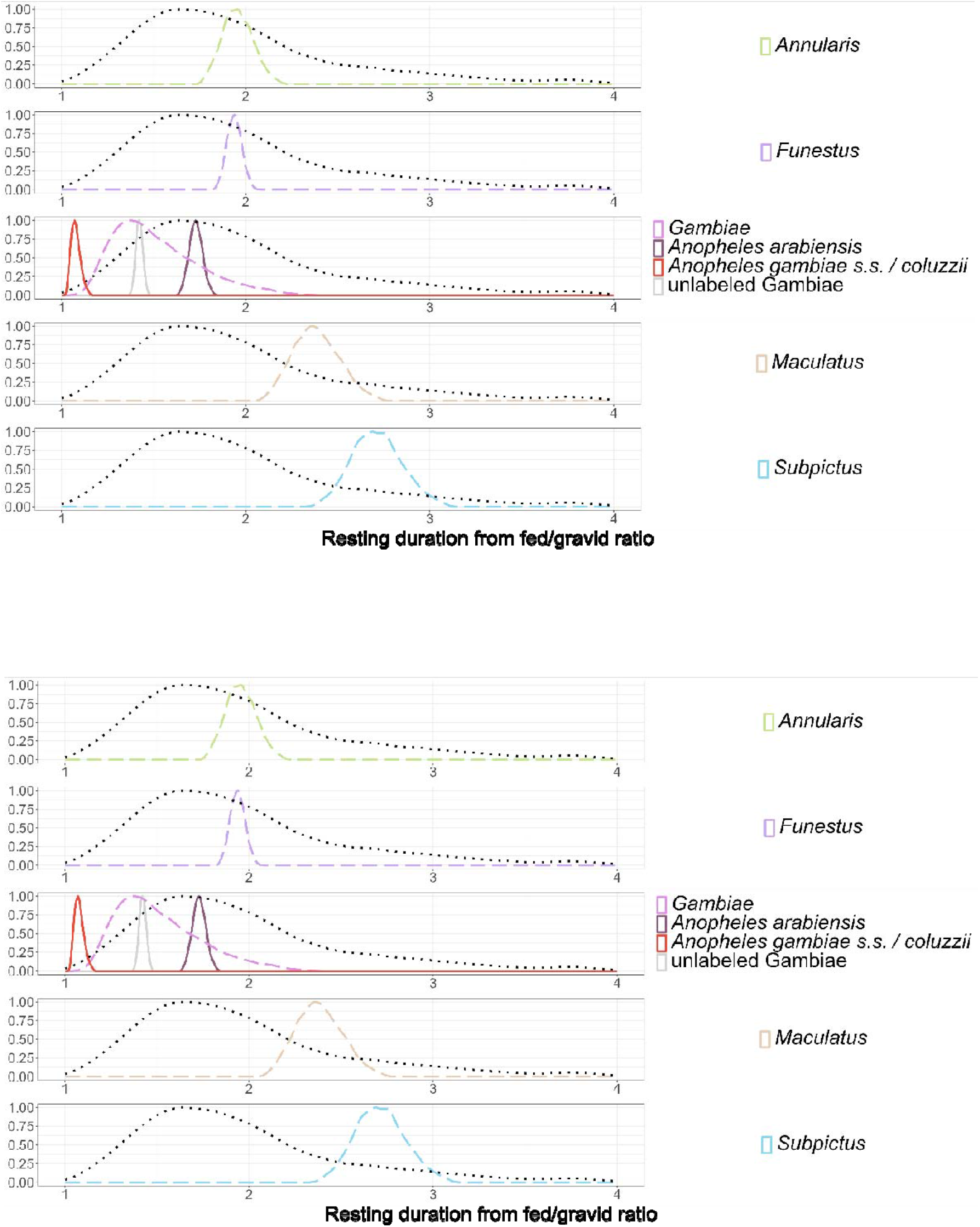
Posterior densities for the resting duration. The black dotted line is the posterior density for the pooled estimate, the coloured dashed lines, for each complex, and the coloured solid lines for individual species. Y-axis is normalised to ease visualisation of all curves.

### Sac rate

The raw weighted average for the sac rate was 47 ± 29% and the Bayesian estimate for the whole *Anopheles* genus was 48% (21-76) (cf. Supplementary Figures 18 and 19 for further examples on the *Gambiae* and *Funestus* complexes/groups). *Anopheles albimanus, Funestus* group and *Gambiae* complex were the only names with sample sizes larger than 500 mosquitoes and their posterior densities are also the most peaked, with mean estimates of respectively 44% (30-58), 52% (39-63) and 56% (43-69) (Figure 10).

**Figure 10.**
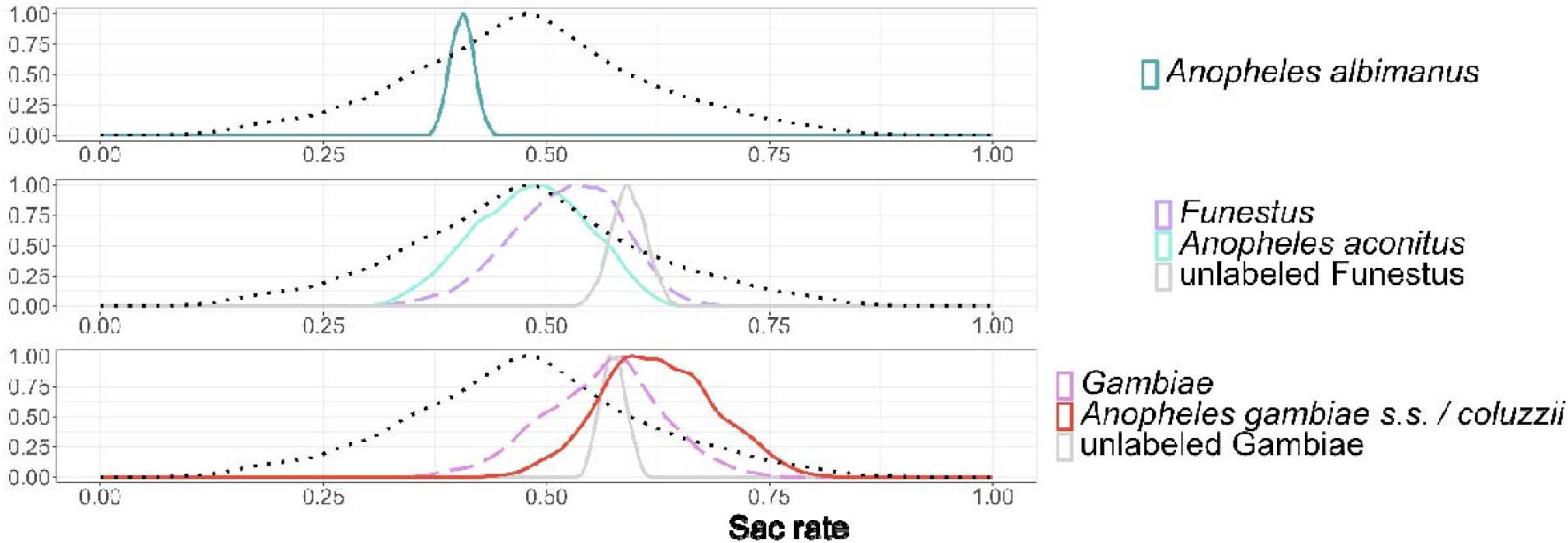
Posterior densities for sac rate for a subset of species. The black dotted line is the posterior density for the pooled estimate, the coloured dashed lines, for each complex, and the coloured solid lines for individual species. Y-axis is normalised to ease visualisation of all curves. The full figure including all species is provided in the Supplementary material (Supplementary Figure 15).

**Figure 11.**
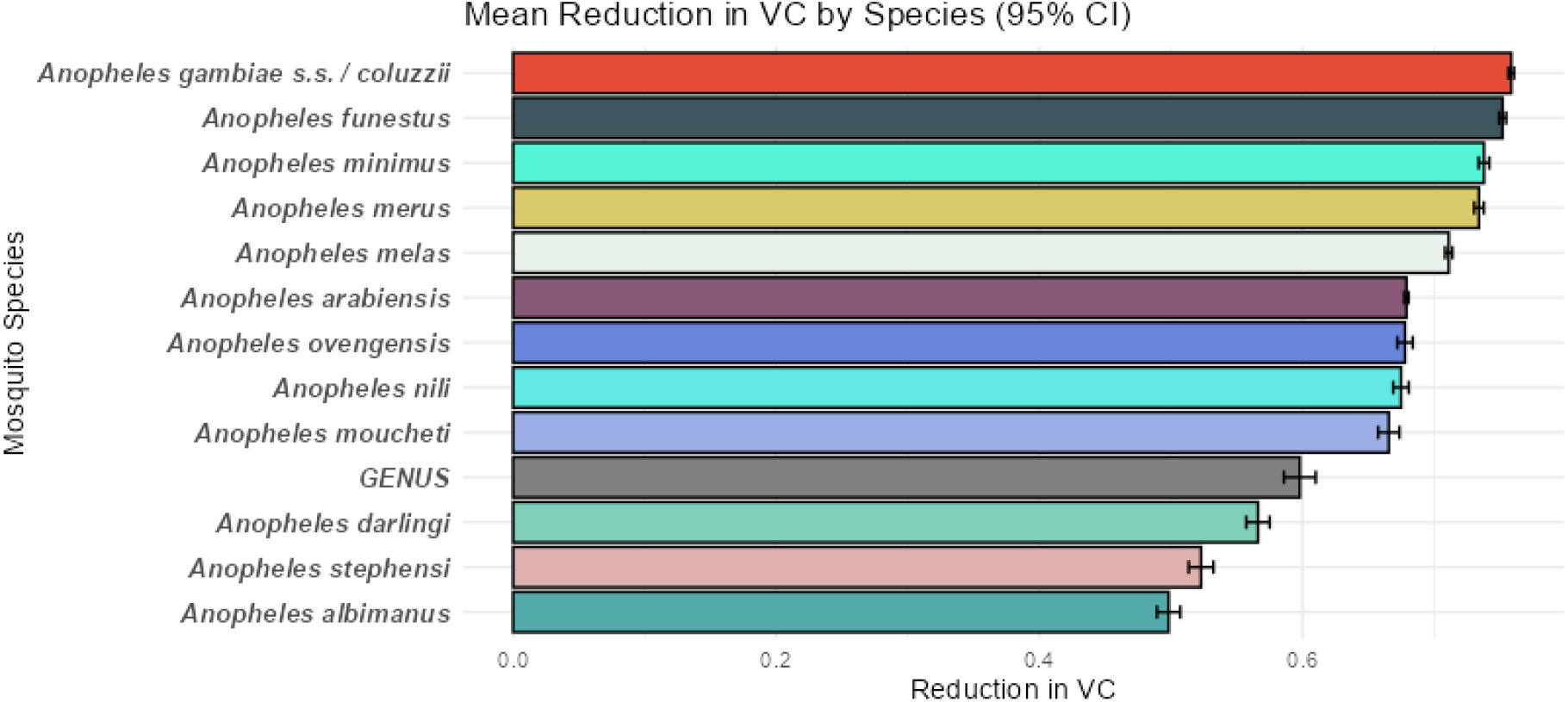
Average reduction in vectorial capacity (VC) following an 80% coverage with the chlorfenapyr-alphacypermethrin ITNs for some mosquito species with corresponding 95% credible intervals. The full table of mean VC values and variability measures for all species is provided in the Supplementary material (Supplementary figure 18).

### Reduction in vectorial capacity

We observe that the average reduction in vectorial capacity (VC) following an 80% coverage with the chlorfenapyr-alphcypermethrin nets varies considerably across mosquito species (Figure 11). *Anopheles gambiae s*.*s*. / *coluzzii* shows the highest mean reduction in VC (75.9%, 75.7-76.1), followed by *Anopheles funestus* (75.2.%, 74.9-75.5) and *Anopheles minimus* (73.8%, 73.4-74.2) In contrast, *Anopheles albimanus* exhibits one of the lowest average reduction (49.8%, 48.9-50.7), while the genus-level estimate remains relatively moderate (59.8%, 58.6-61.0).

Regarding uncertainty, the 95% confidence intervals indicate that the degree of variability differs across species. For example, *Anopheles arabiensis* displays a relatively narrow interval, with VC reduction values ranging from 67.7% to 68.1%. Conversely, *Anopheles moucheti* shows a wider interval (65.8% to 67.4%), reflecting higher variability (Figure 11).

When analysing the Partial Rank Correlation Coefficients (PRCCs) for the reduction in VC after the deployment of chlorfenapyr-alphacypermethrin ITNs, indoor HBI shows the strongest positive correlation (0.79), followed by sac rate (0.62) and parous rate (0.57). Endophagy shows a very weak effect (0.00). This indicates that intervention effectiveness is expected to be higher on vectors with a longer life expectancy and that bite preferentially humans and indoors. Resting duration, in contrast, is negatively correlated with vectorial capacity, with a PRCC of -0.23 (Supplementary figure 19).

## Discussion

We developed a statistical model to estimate different bionomics and behavioural parameters for a wide range of *Anopheles* species and complexes from the data in (Massey et al., 2016). Thanks to this information, we were able to quantify for each species the expected vectorial capacity reduction after the introduction of a pyrethroid-pyrrole ITN. This work shows how we can leverage available data to generate species-specific estimates of bionomic parameters to inform local characteristics and behaviour of malaria vectors.

With our new methodology it is now straightforward to generate appropriate parameterisations for mathematical models to represent any site where the vectors are known. In line with previous literature (Briët et al., 2019; Golumbeanu et al., 2024) our work highlights how the effectiveness of vector control interventions such as ITNs can vary across *Anopheles* species, especially because of differences in human blood index, sac rate and parous rate (Wang et al., 2024), as well as variation in human and mosquito activity patterns (Golumbeanu et al., 2024). While we presented only one intervention as an example, a pyrethroid-pyrrole ITN as evaluated in one experimental hut trial (Odufuwa, in prep), the estimates could easily be computed for a wide range of other vector control tools evaluated in experimental hut trials (Champagne et al., 2025; Denz et al., 2021; Fairbanks et al., 2024; Golumbeanu et al., 2024). Such species-specific model parameterisations can therefore be used to increase realism in mathematical models used for subnational tailoring of malaria interventions in various geographies (World Health Organization, 2025).

The method provides both point estimates that can be applied for each vector identified in an area, and measures of uncertainty around these, appropriately estimated using a Bayesian hierarchical model. The Bayesian approach provides a systematic and transparent methodology for weighting observations within and between species, not only ensuring that each study’s sample size is appropriately accounted for, but also using additional information supplied by related surveys to derive shrinkage estimators. These estimates will generally be more accurate than rates and proportions directly estimated from local data which can be highly inaccurate because of sampling variation. The related surveys include both those that sampled the same taxon in a different place, and also in the case of species with few observations, surveys of related taxa, to the extent that the bionomic traits correlate with the taxonomy. This approach even enables parameter estimation for species or complexes with no direct observations, by borrowing strength from related taxa. This is particularly important given the limited data available for some parameters and taxa. The credible intervals provide a measure of uncertainty which users can use to assess the robustness of the estimates.

We could compare our quantitative estimates to the qualitative categorisations of species in the literature. We found, for example, that 2% (0-7) of *Anopheles darlingi* were endophilic, and *Anopheles darlingi* has been reported as exophilic (Marianne E. Sinka, Rubio-Palis, et al., 2010). We also found that only 36% (35-38) of *Anopheles farauti* are endophagic, and the species is indeed reported as exophagic (Sinka, Bangs, et al., 2011). However, comparisons for endophagy should be performed carefully as the types of studies selected (human landing catches) do not capture the opportunistic behaviours of mosquitoes, since collectors are both indoors and outdoors. The estimate of endophagy used here measures the preference of the mosquitoes for feeding indoors, in the presence of available hosts both indoors and outdoors. In real-life settings, where most people are sleeping indoors at night, the proportion of feeding indoors will be different from the endophagy estimated from paired (indoor and outdoor) human landing collections (Monroe et al., 2020; Rozi et al., 2025). As this behaviour parameter is dependent on the location of humans and the relative abundance of vectors’ preferred hosts, methods performing weighting by host location (Golumbeanu et al., 2024; Monroe et al., 2020) can be used to allow for these effects in estimating the true proportion feeding indoors. In this work, we report endophagy as the unweighted outcome of paired indoor-outdoor HLC, but incorporate weighting by host location as part of the vectorial capacity modelling, following (Golumbeanu et al., 2024). In addition, interventions with deterrent effects also modify the availability of hosts and hence indoor feeding: in this work, we estimated endophagy in the absence of vector control, and allowed for modification of host availability as part of the vectorial capacity modelling (Chitnis et al., 2008).

The study-selection process highlighted the lack of data available for some collection methods. This is most visible with endophily, for which we only selected four studies resulting in eight observations on two complexes and one species. Additionally, indoor and outdoor resting can be challenging to quantify appropriately with PSC and exit traps, as studies have observed cow-fed *An. arabiensis* resting in experimental huts (Kitau et al., 2014). Endophily can also be measured with aspiration and other works have highlighted the lack of standardization in measures of resting behaviour (Irish et al., 2024). There were also no data in the (Massey et al., 2016) database on the sac rate, which is included as a parameter in the mosquito feeding cycle model of (Chitnis et al., 2008). Additionally, we found that HBI values strongly varied depending on the location of collection, except for some species showing strong anthropophilic behaviours. Overall, the collection of bionomic data has not generally been done using standardized protocols in all locations, which creates challenges not only for modelling, but for general understanding of mosquito bionomics. There is a great need for improved methods of measuring bionomic data, as well as standardization of protocols in all locations (Malaria Elimination Initiative, 2020).

Our results can be directly reproduced and expanded thanks to the `AnophelesBionomics` package (https://github.com/SwissTPH/AnophelesBionomics). This can be used to explore all the results dynamically by visualising inputs and outputs across species and complexes. The package allows users to run the analysis on their own data, so updated analyses can be run based on more recent local data. Filtering by geographic region and time period means the analysis can be reproduced focusing only on a specific subset of the entire dataset, hence providing estimates that are more locally informed and conversely less driven by the phylogeny. The possibilities of exploring variation in space and time could be exploited to study determinants of bionomic traits that are not currently in the model, for instance environmental variables, or malaria intervention coverage data.

The model only attributes a limited proportion of interspecific variation to the phylogenetic information. Bionomic characteristics may evolve quickly (Sougoufara, Diédhiou, Doucouré, et al., 2014) and may therefore not strictly align with taxonomic relationships. In principle, local selection pressures (for instance due to vector control) can lead to convergence in the characteristics of the different vectors, but it could also lead to divergence if competitive exclusion applies to their distinct ecological niches. Local ecological factors, such as the type of housing or climatic factors, may also influence species behaviour and traits beyond phylogeny. Nonetheless, the use of taxonomy allowed us to use data on complexes and related species to make estimates when the data are very sparse.

This analysis has a number of limitations. Firstly, we had to make some simplifying assumptions in the structure of our hierarchical model. We only included three levels in the hierarchy as including more levels for groups and subgroups would have massively increased the number of parameters to estimate. The current model also assumes all observations to be independent, even if they were from the same study.

Secondly, our species classification relies on the availability of data in the published literature. When classifying the names in the (Massey et al., 2016) database, we always assumed the species had been correctly identified, which is certainly not the case as challenges of mosquito identification have been noted elsewhere (eg. (Rahola et al., 2022)). Additionally, there remains considerable molecular and morphological work to ensure that *Anopheles* systematics are up to date. Using geographical distribution information could allow for more precision in the species classification: *Anopheles minimus* and *Anopheles funestus* both belong to the *Funestus* group, but the first is found in Asia while the second is a dominant vector species of Africa. This could allow us to attribute all the *Funestus* mosquitoes collected in Asia to the *Anopheles minimus* species. Our categories could also evolve if more names were added to the database. For example, the *Pseudopunctipennis* complex could have its own category if some species of this complex were sampled, and the *Hyrcanus* group would separate into *Lesteri* subgroup and *Sinensis* complex if they were sampled.

Thirdly, our quantification of vectorial capacity reduction did not account for species-specific differences in resistance levels. This assumption was made because all current vectors are considered susceptible to chlorfenapyr (Messenger et al., 2023; Sovi et al., 2024). We also assumed that differences in response to pyrethroid-pyrrole ITNs were attributed to differences in bionomics parameters only, despite recent evidence that mosquito species may respond differently to insecticides (Oruni et al., 2026). Accounting for such differences is beyond the scope of the current work but should be explored in future research.

Finally, the database (Massey et al., 2016) only includes data published before 2010. There is an urgent need to update the database with recent data, such as those identified by (Wang et al., 2024), to increase the number of relevant studies and sampled species, and thus of observations. Mass net distributions also started at the beginning of the years 2010s (Global Malaria Programme, 2007), so collections after 2010 would account for the modifications of *Anopheles* behaviour due to vector control interventions (Sherrard-Smith et al., 2019b), as well as changes in human behaviours (Esayas et al., 2024; Monroe et al., 2019; Namango et al., 2024; Odero et al., 2024), climate change (World Health Organization, 2022) and invasive species (Ahmed et al., 2022; World Health Organization, 2022) . `AnophelesBionomics` allows for the integration of new data, facilitating future updates of the database.

In conclusion, we provide a framework to estimate species-specific *Anopheles* bionomic parameters, by summarising available entomological evidence and borrowing information from neighbouring species. Our estimates can be used as indicators of local *Anopheles* species bionomics and behaviour to guide vector control decisions. Our framework leverages existing data to maximise its use both by modellers and decision-makers, and can be reused with newly generated data to keep updating knowledge on *Anopheles* bionomics.

## Author contributions

Conceptualization: TAS. Identification of collection methods: BZ. Selection of the taxonomy: BZ. Implementation of the statistical model: JL, AT. Software: AT. Analysis of the mathematical model: AT, CC. Resources: OGO, SM. Supervision: CC, EP. Funding acquisition: EP. Writing, first draft: JL. Writing, review and editing: JL, AT, TAS, BZ, MG, OGO, SI, SM, EP, CC.

## Code availability

All code and data are available at https://github.com/SwissTPH/AnophelesBionomics.

## Acknowledgements

This work was supported in whole or in part by the Gates Foundation [INV-068864]. The conclusions and opinions expressed in this work are those of the author(s) alone and shall not be attributed to the Foundation. **Under the grant conditions of the Foundation, a Creative Commons Attribution 4.0 License has already been assigned to the Author Accepted Manuscript version that might arise from this submission**.

The funders did not play any role in the study design, data collection and analysis, decision to publish, or preparation of the manuscript.

This work also received financial support from the Geigy Foundation.

